# Automated and high throughput measurement of leaf stomatal traits in canola

**DOI:** 10.1101/2024.06.12.598768

**Authors:** Lingtian Yao, Susanne von Caemmerer, Florence R Danila

## Abstract

**Background:** Automating stomatal trait measurement has gained popularity because of their inherent importance for field phenotyping application as stomata are critical for both carbon capture and water use efficiency in plants. Such tool has been reported for rice, wheat, tomato, barley and oil palm. However, none exist yet for canola, which is an important economic and agronomic crop globally.

**Results:** We developed a new toolkit called Stomatal Comprehensive Automated Neural Network or SCAN by combining the use of high-resolution portable digital microscopy with machine learning based on You Only Look Once algorithm (YOLOv8). Digital micrographs of leaf surfaces enter the SCAN pipeline, which includes stomata detection, stomata segmentation and stomatal pore segmentation models, to measure stomatal density, stomatal size and stomatal pore area, respectively. In addition to SCAN’s ability to measure leaf stomatal traits in canola at 89 to 94% accuracy, we also showed that SCAN can be used to predict stomatal density even in species not included in the training set such as Arabidopsis, tobacco, rice, wheat, maize and proso millet. SCAN was designed for the biological science community with the premise that users are not required to possess advanced programming capabilities to manage dependency prerequisites, execute the models, and integrate the analysis. This was achieved by packaging the models into a desktop application system that can be accessed offline.

**Conclusion:** Overall, SCAN provides a non-destructive, real-time, portable, and high-throughput measurement of leaf stomatal traits in canola. The minimised hardware requirement and user-friendly desktop application system make SCAN suitable for field phenotyping application.

## Background

Canola (*Brassica napus* L.) production has economic and agronomic advantages. It is grown as one of the most important oil seeds, accounting for the world’s second vegetable oil production after soybean. Global canola production for 2023-2024 is projected at 87.4 million tonnes with top canola producers including Canada, the European Union, China, India, and Australia (Reidy, 2023). The growing demand for renewable diesel is pushing the global canola industry to produce more each year. Canola is well suited for biofuels production because of its high oil content, providing more feedstock for fuel production and less byproduct, and low levels of saturated fat, which is linked to improved cold weather performance (Sey et al., 2020). However, with climate change causing more frequent heat waves and severe drought, canola production and quality are under threat. Hence, there is a need to selectively breed for climate-resilient and more water-use efficient varieties.

The most direct measurement of drought tolerance is based on water-use efficiency and yield under water-limiting conditions (Tuberosa, 2012). Measuring leaf stomatal density can give an indication of plant’s water-use efficiency (Bertolino et al., 2019; Xu & Zhou, 2008) and hence drought tolerance. Biologically, stomata are important “gates” for both gas exchange (CO2) of the leaf during photosynthesis and water vapour during transpiration (Lawson & Matthews, 2020). Since each stoma is made up of two guard cells, which bordered the pore, understanding stomatal traits such as size and pore dynamics are also important (Lawson & Matthews, 2020). While stomatal density can be obtained by simply imaging the leaf under a microscope, this involved cutting the leaf or leaf tissue from the plant (Sai et al., 2023). Meanwhile, the traditional non-destructive way of quantifying leaf stomatal traits using leaf imprint is laborious and time-consuming, and often lab-based due to dependency to conventional microscopes (Kwong et al., 2021; Millstead et al., 2020). As most crop science research head towards the industry direction, translating non-destructive method of stomatal capture for field application is imperative. This means creating a high-throughput approach on image capture but also trait measurement.

Automation of stomatal trait measurement using machine learning-based model has already been reported for other industrially important crops such as rice, wheat, tomato (Pathoumthong et al., 2023), barley (Sai et al., 2023) and oil palm (Kwong et al., 2021). However, to date, none exist for canola. Here, we established a user-friendly toolkit for stomatal trait measurement in canola leaf based on an efficient and high-throughput computer vision inferencing method. This method involves four processes including (1) microscopy image collection and file structuring, (2) models training (3) statistical analysis after model prediction and (4) high throughput system. We called this toolkit Stomatal Comprehensive Automated Neural Network or SCAN. SCAN makes use of portable digital microscope giving researchers a non-destructive and real-time approach to capture stomatal traits in high throughput, while the subsequent inferences and post-processing analysis are automated.

## Materials and methods

### Image capture

Canola (*Brassica napus* L.) plants growing in the glasshouse and field were used in this study. We used a portable digital microscope with 5.0-megapixel sensor resolution and 700x-900x magnification (Dino-Lite Edge 3.0, Cat no. AM73915MT8) to capture the stomata from the top and bottom surfaces of the canola leaves (Fig. 1A). Micrographs were captured at a focused magnification of approximately 691x, with a resolution of 2560 x 1920 pixels and an embedded scale bar of 0.05 mm.

**Figure 1.**
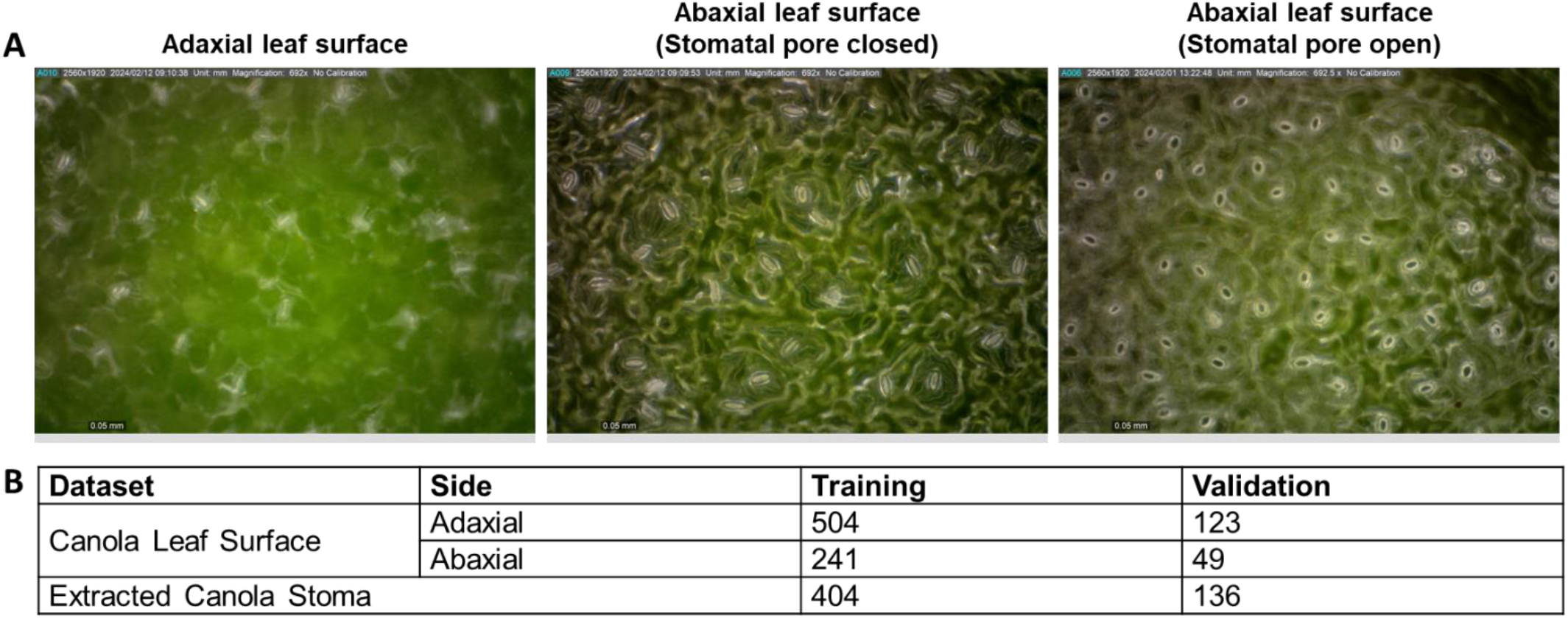
Dataset used for model development in SCAN. Dataset used for the model development in canola include micrographs taken from adaxial and abaxial leaf surfaces capturing variation in stomatal pore status (**A**). Summary of datasets splits in model development is tabulated in **B**. The canola leaf surface dataset is used to train stomata detection and segmentation models. The extracted canola stoma dataset is generated from stomata detection model and is used to train the pore segmentation model.

### Preprocessing

The labelled inputs are required on any supervised learning structure to identify the task before performing a function mapping to deliver the desired prediction (Nasteski, 2017). Leaf micrographs from canola were manually annotated for stomatal traits such as stomatal density, stomatal size and pore area using Roboflow (Dwyer, 2024). We used a polygon to outline each stoma (i.e., area occupied by the stomatal pore and corresponding guard cells) to generate our dataset for both stomatal detection and segmentation purposes. We selected distinguishing stomata with different degrees of aperture status from stomatal detection prediction results and applied the same annotating strategies to build the dataset for pore segmentation. These datasets were split into training and validation sets ratios of 0.8/0.2 and 0.75/0.25, respectively (Fig. 1B).

### Model training

The entire detection and segmentation tasks were carried out by using either You Only Look Once (YOLO) algorithm modular templates or transferred learning from our pretrained model. The workflow is composed with two tasks in each stage of model training: (1) initializing YOLO architecture template from scratch to classify and localise each stoma region of interest (ROI), enabling transferred learning for stomata size and pore segmentation, and (2) fine-tuning the hyperparameters to improve models’ performance.

As depicted in Figure S1, YOLOv8 consisting of two modules was used. These modules are (a) CSPDarkNet53 (Wang et al., 2020) feature extractor in the backbone including Convolution(conv), Cross Stage Partial bottleneck with 2 convolutions - Fast (C2f) and Spatial Pyramid Pooling - Fast (SPPF) units, and (b) the head equipped with Path Aggregation Feature Pyramid Networks (PAFPN), which fuses multi-level features followed by the final prediction layer to deliver detection or segmentation tasks. The purpose of the training in deep learning theory is to find a best non-linear transformation between input and output through the interaction of the error evaluation in output layer and backpropagation updating mechanism (LeCun et al., 2015). To find best fitting pattern, the adjustable parameters (so called weights) were optimised.

All original images in datasets with 2560 x 1920 resolution pixels were resized to 640 x 480 with RGB channels during the data pre-processing stage. This resizing was done to avoid model over-complexity and increase the training speed. The stomatal leaf surface dataset was initially used to train the stomata detection model with the default hyperparameters setting in YOLOv8 for 200 epochs. However, two major obstacles were encountered: [1] incomplete or cropped stomata at the edge of the microscope images and [2] blurred or out-of-focus stomata. To address these issues, we employed K-folder cross-validation strategy (Mosteller & Tukey, 1968) and a higher training image resolution of 1024 * 792 were employed to enhance model learning without requiring an extended dataset and additional annotations. As a result, the detection model was able to classify the two types of stomata described in Figure 2. The stomatal dataset was split into K partitions with each partition used as a validation set to evaluate the model. Subsequently, the overall average performance across the K partitions was recorded for model evaluation.

**Figure 2.**
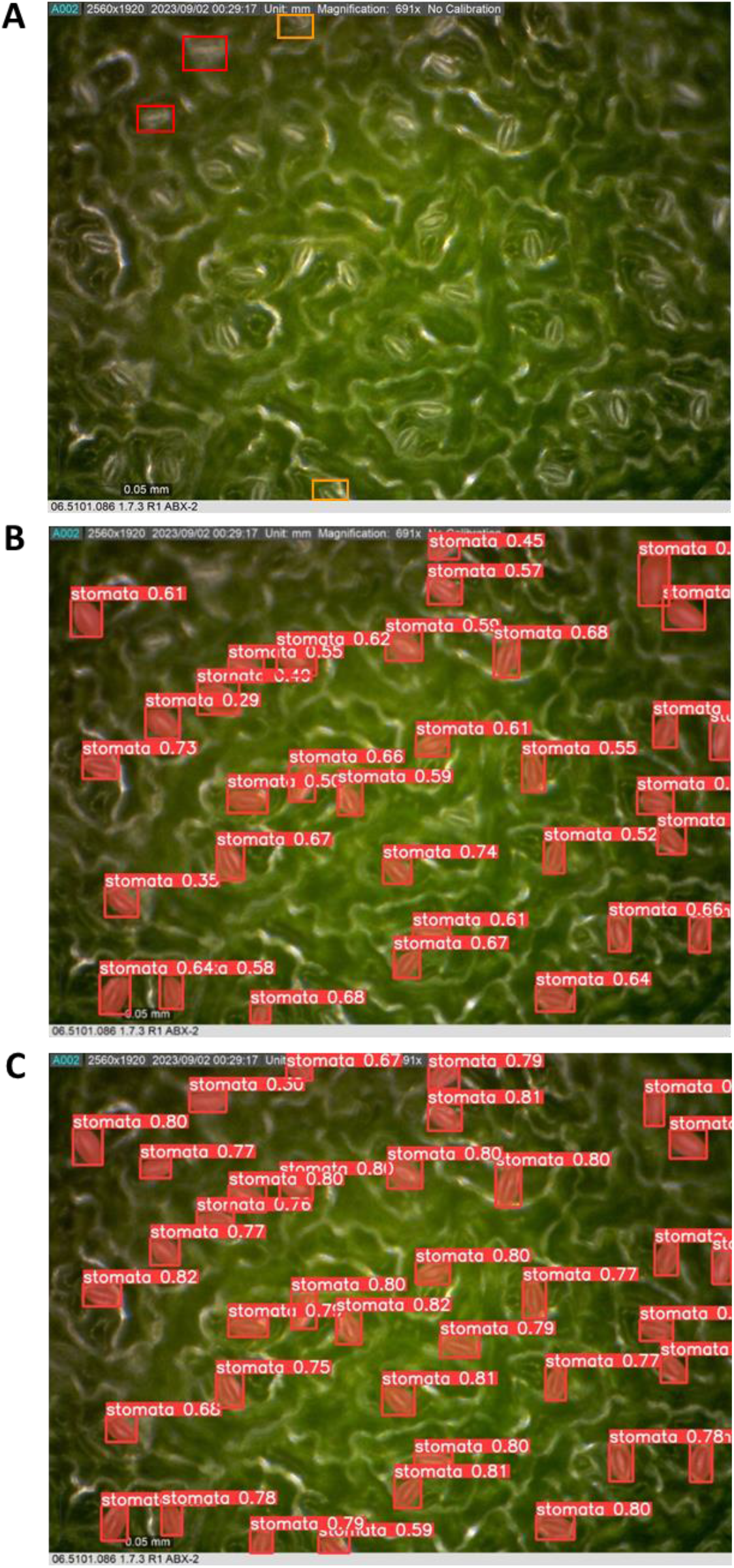
Detection of out-of-focus and incomplete stoma in the image using SCAN. In **A**, orange boxes indicate examples of incomplete stomata, while red boxes highlight out-of-focus stomata captured during imaging. **B** shows that the model has limited ability to detect these features. The confidence values, indicated by the numbers in the image, were <0.70 for most of the detected stomata. However, by increasing the input image dimensions from 640 x 480 to 1024 * 792 and employing K-folder cross validation strategies, the model’s ability to correctly detect and localize these features is enhanced as shown by the increased number of annotated stomata and greater confidence values (**C**).

After several training evaluations of the detection model, the following hyperparameters fine-tuning strategies were carried out to improve the model performance. Weights regularization was adjusted using the AdamW optimizer (Loshchilov & Hutter, 2017) with a learning rate range from 1E-3 to 1E-6, in addition to applying dropout (Srivastava et al., 2014) and mosaic data augmentation to enhance model generalization on unseen datasets. With the optimal results from fine-tuning for detecting and localizing each stoma in the microscope images, the weights from detection model were transferred to train the stomatal segmentation model. Concurrently, regions of each representative stoma with either open or closed pores were cropped to establish a pore dataset. The pore segmentation model applied YOLOv8-nano structure templates with transferred weights from the stomatal segmentation model, trained and fine-tuned using the domain knowledge of the extracted pore dataset. The details of the three model training templates and transferred weights are summarized in Table S1.

The training tasks were carried out on an Ubuntu 20.04 Linux server at the Research School of Biology in Australian National University, using two Nvidia A30 (24G) Graphic Processing Units (GPUs). The full details of models’ weights, hyperparameters, training scripts and datasets can be found at https://github.com/William-Yao0993/FD_detection.

### Model evaluation

Mean Average Precision (mAP, Fig. 3) and F1 score were used to assess model ability (Fig. 4). mAP is calculated as the mean value of each class area under the precision-recall curve over thresholds, and the F1 score is the harmonic mean of precision and recall. The formulas are defined as follows:

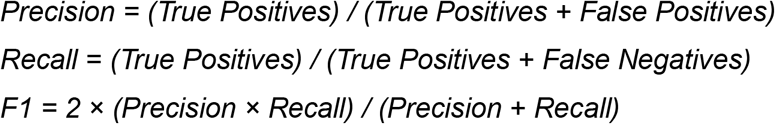

**Figure 3.**
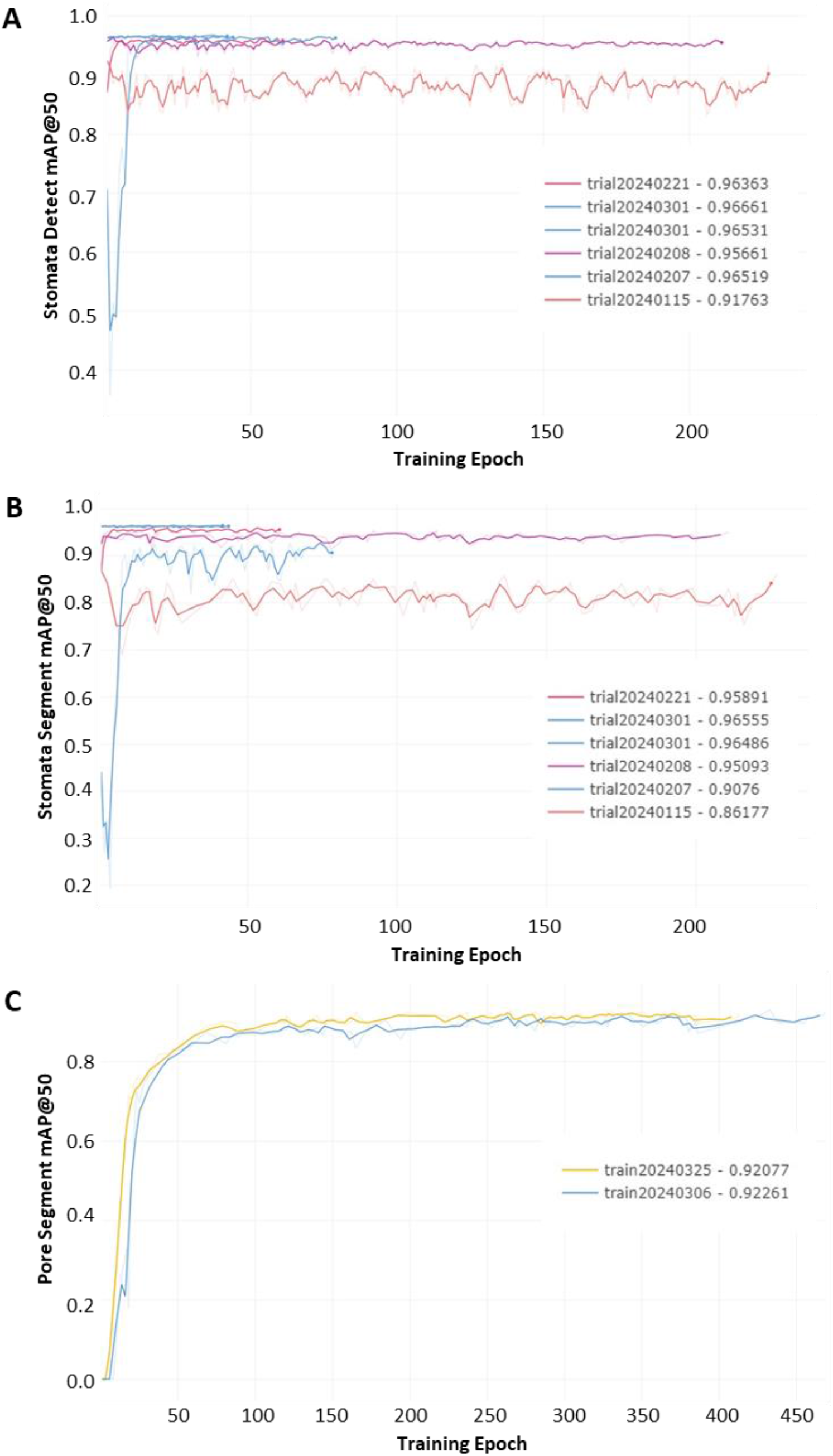
Mean average precision over 0.5 confidence-level (mAP@50) in training stage of the three models in SCAN. The trends in mAP@50 during the training stages for stomata detection (**A**), stomata segmentation (**B**), and pore segmentation (**C**) models show how the models’ performance improved and kept the peak over time.

**Figure 4.**
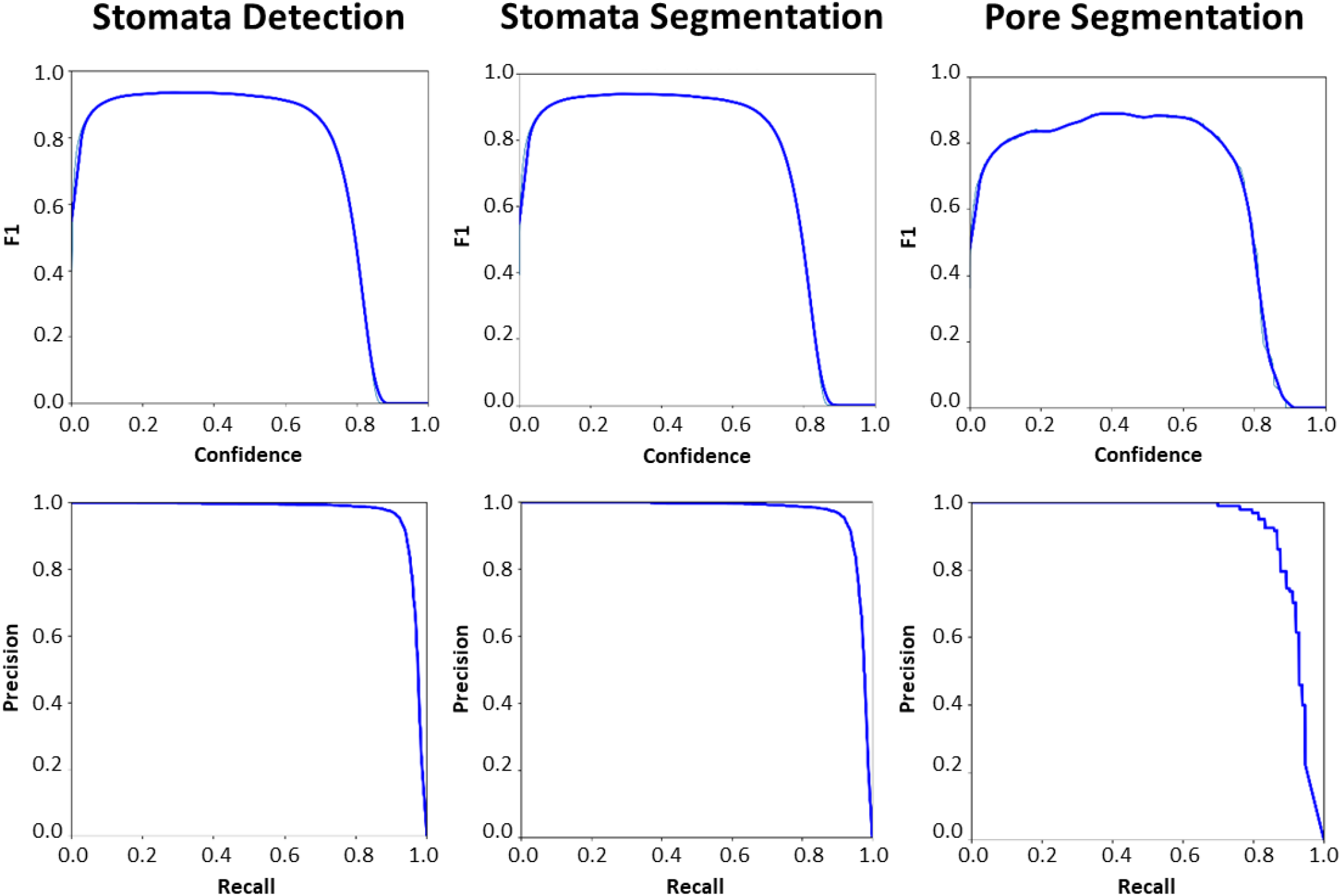
F1-Confidence and Precision-Recall curves of all fine-tuned models’ performance in SCAN. The performance of all fine-tuned models in SCAN, including stomata detection, stomata segmentation, and pore segmentation was evaluated. The F1-Confidence curves (top panels) show the relationship between the F1 score and the confidence threshold for each model. The F1 score is the harmonic mean of precision and recall, and these curves help to determine the optimal confidence threshold for the best performance. The F1 score remains high across a wide range of confidence levels in both stomata detection and segmentation, indicating the robust performance of these models. The F1 score of pore segmentation is stable but relatively less precise as the other models. The Precision-Recall (PR) curves (bottom panels) illustrate the trade-off between precision and recall for each model. High precision and recall values indicate effective model performance. Like F1-Confidence curves results, the PR curves show both high precision and recall in stomata detection and segmentation models, while pore segmentation model has a slightly drop in recall at higher precision levels.

### Scale bar detection

The results from model inferences, including both bounding boxes from detection and masks from instance segmentation, were at the pixel level. These measurements cannot directly correlate in the bioinformatic domain without standardization to real-world units. To address this, an automated algorithm was applied to detect the scale bar in the microscope images and return a pixel-to-unit ratio. A combination of Canny edge detection (Canny, 1986) and Hough line transformation (Duda & Hart, 1972) was used to identify line candidates in the image, in this case, the scale bar was embedded at the bottom-left of the input image. Line candidates were filtered based on rules in polar coordinate system, i.e., have angles in polar coordinates near 0 or 90 degrees, with a tolerance of ± 2 degrees. The line with the maximum length was chosen as the result. The ratio was calculated between scale bar unit and the detected line. In cases where no lines fit the rules-based algorithm, a default ratio was used. The pseudocode of the scale bar detection algorithm is provided in Figure S2. With the connection between pixel and real-scale units, the following three measurements were derived: stomata density, stomata size and pore area. Stomata density was determined by the numbers of bounding boxes in stomata detection model, while stomata size and pore area were calculated based on the masked area in pixel and converted to real scale using the scale bar ratio from stomatal segmentation and pore segmentation models, respectively. This process addresses the pipeline of SCAN in a single microscopy image, as depicted in Figure 5A.

**Figure 5.**
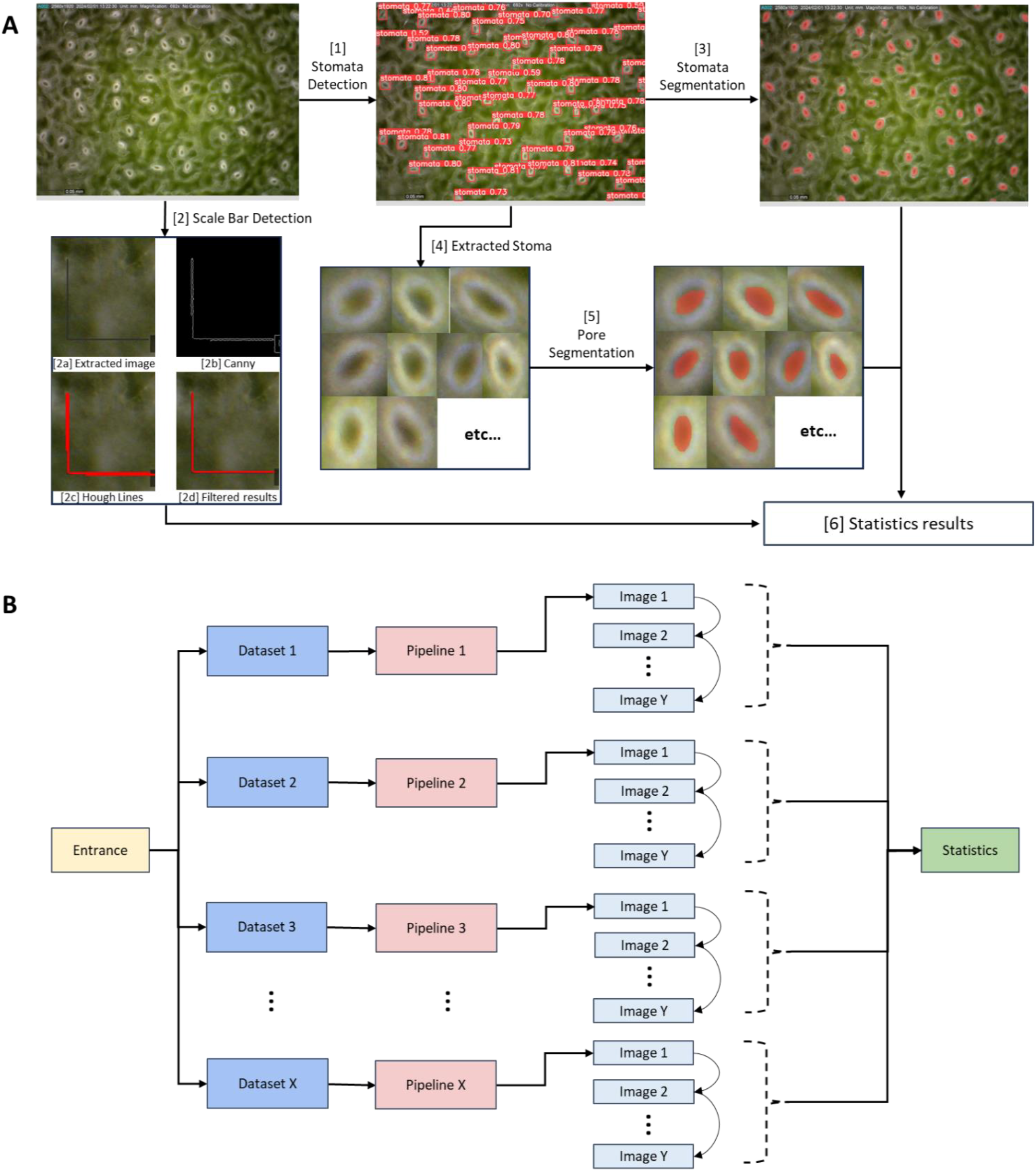
SCAN measurement pipeline and system design. The pipeline of SCAN (**A**) involves several key steps to process and analyse microscopy images for stomatal traits. The input microscopy image is used to detect and localize all possible stomata above a confidence threshold [1] and run the scale bar detection algorithm [2]. The localised stomata from [1] is then used to initiate stomata segmentation model [3] to create precise segmentations in pixel level, and each stoma within the bounding boxes is extracted [4] to form a dataset for pore analysis. The extracted stoma dataset from [4] enter the pore segmentation model [5] to identify and segment the pore areas. The final step integrates all the information from the stomata detection [1], the scale bar detection [2], the stomata segmentation [3], and the pore segmentation [5] to generate comprehensive statistical results [6]. This pipeline ensures a thorough and automated analysis of stomatal traits, from initial detection to detailed segmentation and statistical evaluation. The parallel system design of SCAN (**B**) enables users to provide a main folder containing multiple subfolders as an entrance. SCAN works through the structure from the entry point and enables parallel processing of each subfolder. Within each subfolder, SCAN sequentially processes each image through a single pipeline. All information from the parallel pipelines is then gathered for statistical analysis.

### High-throughput system and application

Beyond the prediction on a single image, a parallel computing system using Central Processing Units (CPUs) in conjunction with the multiple model engines to significantly reduce model inference and statistical analysis time was integrated in SCAN to improve the system performance on standard laboratory workstations that lack a GPU and large random-access memory (RAM). This enables SCAN to perform inferences and computations at multiple folder levels, with each folder containing microscopy images. Here, the input folder was first searched and structured to target image type files (supported formats: jpg, jpeg, png, bmp, dng, mpo, tif, tiff, webp, pfm). The system was designed to automatically determines whether to enable multiple model engines based on the folder structure. For example, if the input folder contains x number of subfolders of images as shown in Figure 5B, the system enables x number of models to run in parallel threads, performing statistical analysis. As results, the system generates an excel file that contains each stoma-level information (stoma size and pore area in mm^2^), each image-level summary (stomata density per mm^2^, statistical mean and standard errors for stoma size and pore area), and each subfolder-level summary (statistical mean and standard errors for stomata density, size, and pore area). Boxplots for these aspects were generated for distribution comparison among folders. Additionally, SCAN was integrated into an interactive and user-friendly application that researchers can interact with, as depicted in Figure 6. The inferences results predicted image with annotations, statistical tables, and plots can be visualized in the application or exported. Detailed application instruction and download information can be found at https://github.com/William-Yao0993/SCAN/blob/main/README.md.

**Figure 6.**
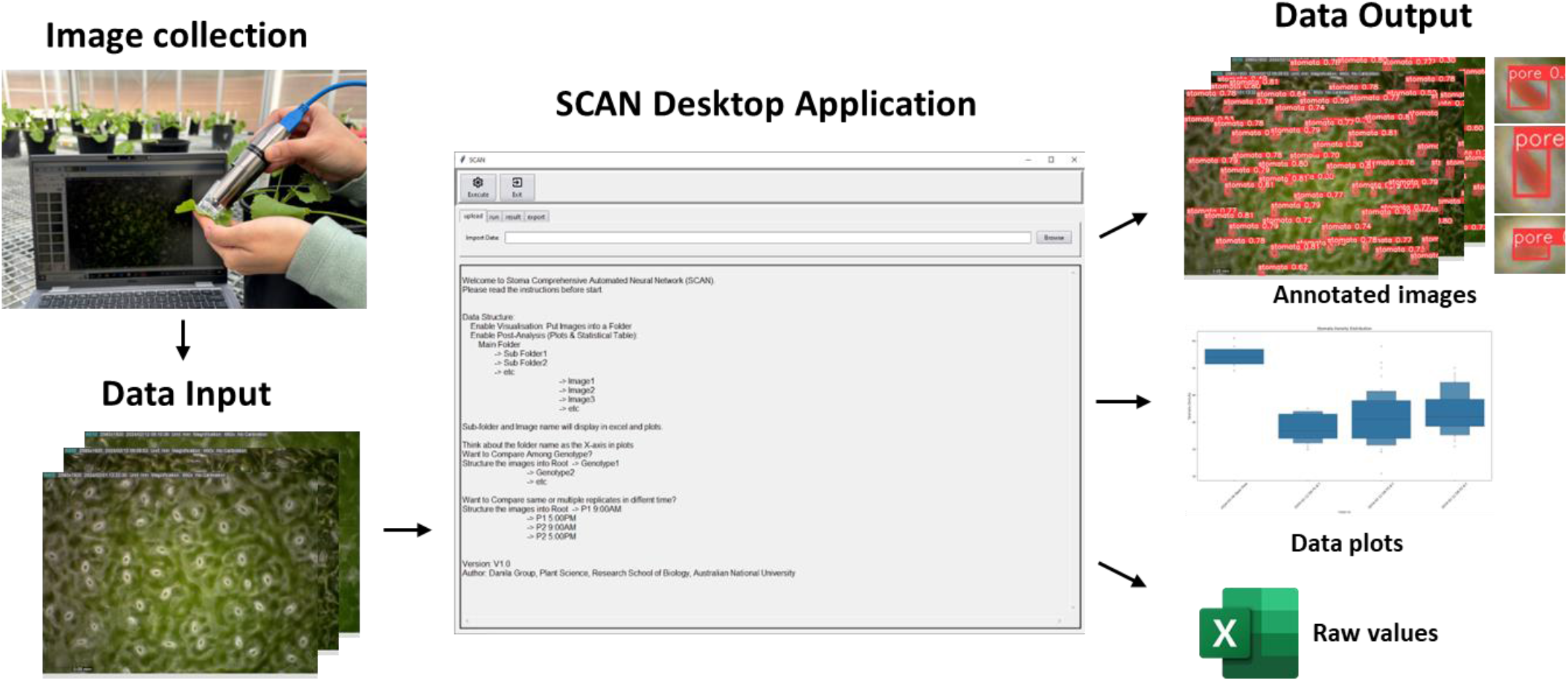
SCAN application workflow. Leaf micrographs obtained from portable digital microscope are uploaded into SCAN, where stomata detection, stomata segmentation and pore segmentation models will be applied simultaneously. Resulting exportable data from SCAN include annotated images, data plots and MS Excel file containing raw values. The annotated images contain original micrographs with bounding boxes to indicate the detected stomata, and masks that outline stoma size. Extracted stoma images with bounding box and mask that outline the pore area are also provided as exportable data output. Data plots include stomatal density, stomata size and pore area based on data input structure. Full demonstration and download of SCAN can be found at: https://github.com/William-Yao0993/SCAN.

### Manual-machine correlation test in canola

Stomatal density, stomatal size and pore area were manually recorded and measured from the micrographs using ImageJ software. Stomata density was defined as the number of stomata per mm^2^ leaf area. Stomatal sizes are measured as the area occupied by stomatal pore and corresponding guard cells in mm^2^. Pore area is captured as the area occupied by the stomatal pore in mm^2^. For stomatal density, 66 unseen microscopy images with varying stomata densities were counted. For stomatal size and pore area, the freehand drawing tool was used on 152 extracted image areas containing both open and closed stomata. The SCAN results and manual ImageJ results were compared by plotting SCAN results on the x-axis and manual ImageJ results on the y-axis, fitting a linear regression line, and calculating the Pearson correlation coefficient (Fig. 7).

**Figure 7.**
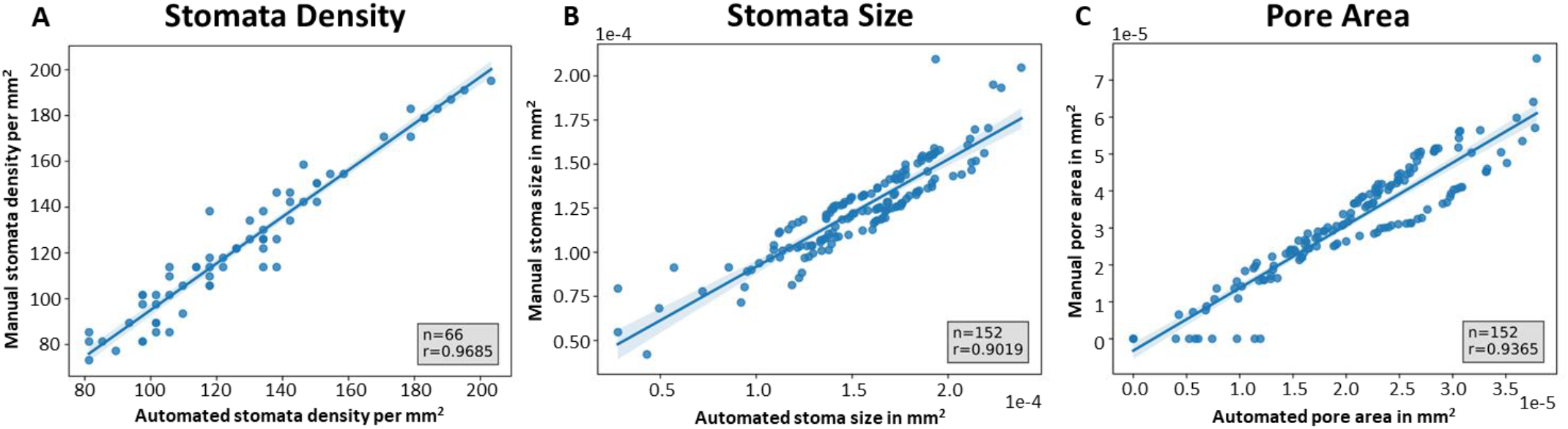
Correlation test between manual and SCAN measurements in canola. SCAN can effectively predict and measure stomata density (**A**), stomata size (**B**) and pore area (**C**) in canola as shown by the high correlation between manual and automated measurements. Each graph plots manual measurements on the y-axis and automated measurements on the x-axis, with a blue line representing the best fit line for the manual-machine results. The Pearson correlation coefficient (R) is indicated in each graph with sample size (n). The test in stomatal density included 66 images while stomata size and pore area used 152 sample stomata.

### SCAN processing time evaluation

The SCAN processing time across three different processing units: a mid-range CPU (Intel(R) Core (TM) i7-6700 CPU @ 3.40GHz 3.41 GHz), a newer CPU (13th Gen Intel(R) Core (TM) i7-1365U 1.80 GHz) and a server GPU (NVIDIA A30 Tensor Core GPU 24G) was compared. Processing time was measured for the entire SCAN system workflow using two test datasets. The 132-image dataset is an unseen dataset containing canola microscopy images of both abaxial and adaxial leaf surfaces, while the 427-image dataset was generated by selecting images from the training dataset. Each microscopy image has resolution of 2560 x 1980. Total images processed and the processing time in seconds for each configuration, both with and without pore measurements were tabulated in Table 1.

**Table 1.**
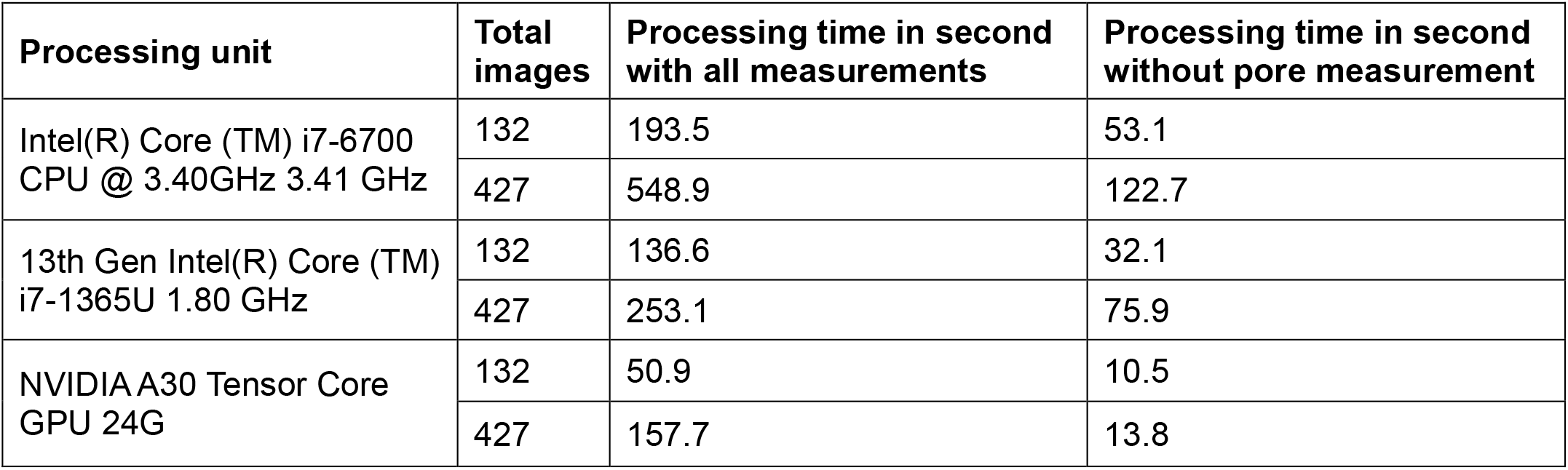
SCAN processing time in different processing units.

### SCAN application to other species

The suitability of SCAN to detect and measure stomatal density was tested in other species. These include two dicot species, *Arabidopsis thaliana* and *Nicotiana benthamiana* (tobacco), and four monocot species, *Oryza sativa* (rice), *Triticum eastivum* (wheat), *Zea mays* (maize) and *Panicum miliaceum* (proso millet). Micrographs were captured at magnification of ∼691x (Dino-Lite Edge 3.0, Cat no. AM73915MT8) and/or ∼414x (Dino-Lite Edge 3.0, Cat no. AM73915MZT4) (Fig. 8A-B). Stomatal densities were quantified manually using ImageJ software and plotted against values obtained from SCAN (Fig. 8C).

**Figure 8.**
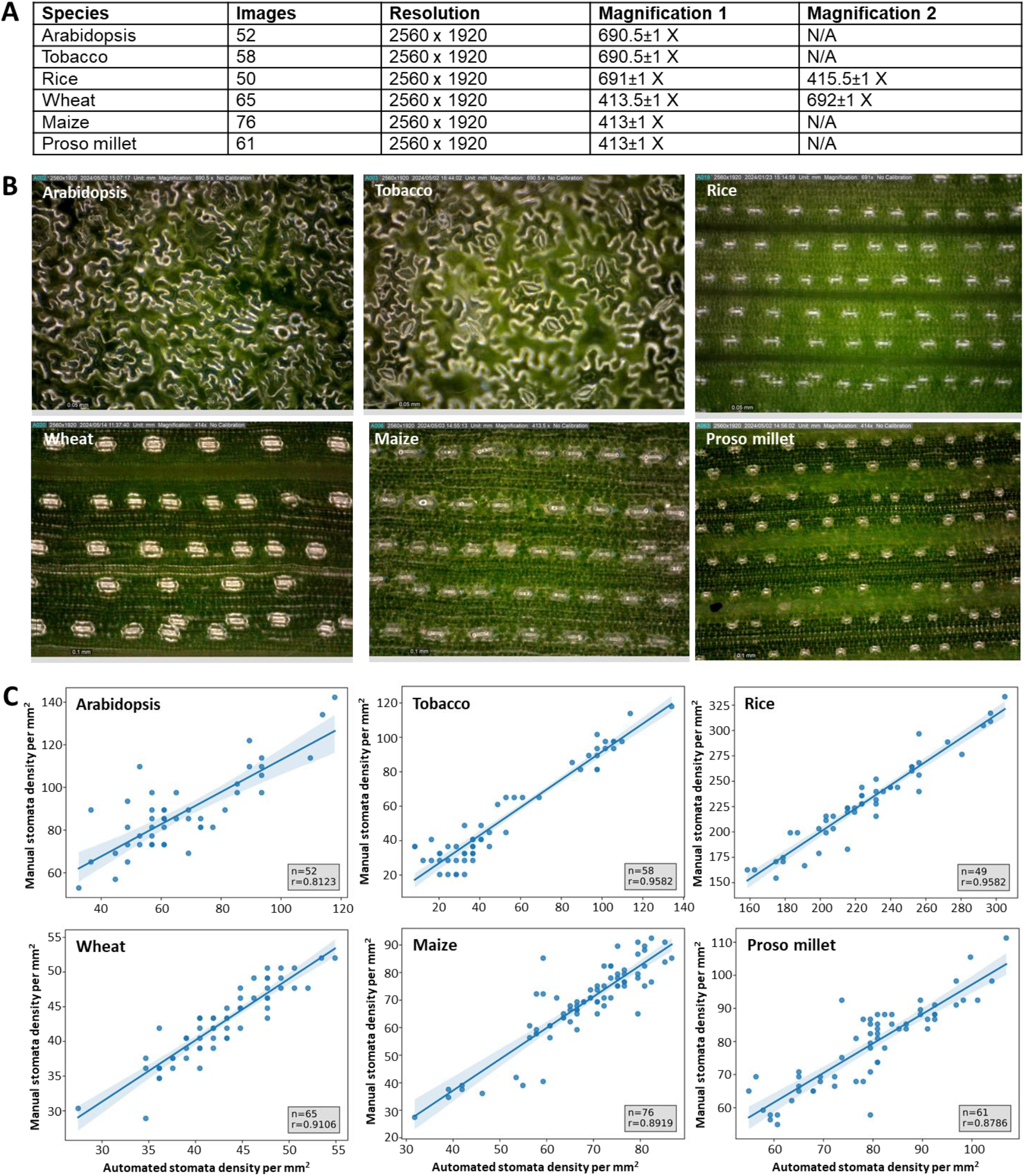
SCAN application to other species. The microscopy image collection setup for Arabidopsis, tobacco, rice, wheat, maize and proso millet is tabulated in **A**. The table includes the number of images, resolution and magnification values used during imaging. Magnification 1 represents the optimal value where the balance between the image quality and stomata capture per image was achieved. Magnification 2 represents backup value that zoom in or out stomata. Representative images obtained from Magnification 1 for the six species are presented in **B**. The comparison between manual and automated stomata density measurements for the six species is shown in **C**. The resulting Pearson correlation coefficients or R, which ranges from 0.8123 to 0.9582, indicates the ability of SCAN to generalise and predict stomata density in both dicot and monocot species that were not part of the training set.

## Results

### SCAN performance in canola

The parameters in each model of SCAN were optimized to deliver the best predicting results for canola. To evaluate the performance of SCAN, we ran machine learning metrices and manual-machine stomata measurement test on leaf images obtained from canola. When tested on the validation set, SCAN achieved the highest mAP at a 0.5 confidence-level (mAP@50) of 0.967, 0.959 and 0.921 (Fig. 3) and F1 scores of 0.94, 0.94 and 0.89 (Fig. 4) for stomata detection, segmentation, and pore segmentation, respectively. Across the three models, the stomatal density and size models performed stably and reliably at the 0.83 confidence level, while the pore area model had slight decreases compared with the other two (Fig. 4). This was not surprising because the input images for the pore segmentation model were the extracted areas from the microscope images which had dimensions of around 80-130 pixels, containing much less information for accurate prediction.

We conducted manual measurement using the ImageJ software to verify the correctness of SCAN for stomatal density, stomatal size and pore area (Fig. 7). We found that the data were highly aligned along the fitted line in stomata density plot (Fig.7A). Estimating stomata area and pore area, however, was more challenging than counting density, especially when measuring pores in nearly closed stomata. The area calculations for stomatal size showed slight differences between manual and SCAN results (Fig. 7B) potentially due to mouse drawing errors and relative inaccuracies. This discrepancy was also observed in pore area measurement, where SCAN tended to predict a small size for nearly closed stomata, while humans judged them as closed (Fig. 7C). Empirically, stomatal pore areas under 1.5E-05 (mm^2^) were determined as closed by human observation, however, there was no significant difference when a threshold was set for recognizing closed and open stomata. Overall, the statistics from SCAN were highly correlated with manual results, achieving correlation coefficients of 0.968, 0.902 and 0.937 for stomata density, size, and pore area measurements, respectively (Fig. 7).

In terms of measurement processing time, SCAN offers significant advantages compared to manual methods or other stomata detection systems. On average, the manual measurement time for one microscopy image ranges from 3 to 6 minutes, depending on the total number of stomata, image quality, and pore aperture opening status. In contrast, SCAN can automate the measurement of each image within 2 seconds. This was tested across three different processing units using two canola testing datasets with varying numbers of images, as summarized in Table 1. The high-throughput performance of SCAN was demonstrated, showing that it can handle hundreds of images within minutes, even with minimal processing units. In addition, SCAN can process and produce results for over 400 images including all measurements, within 157.7 seconds when using a GPU. This automation significantly reduces the workload for researchers, as it would take days to manually measure stomatal density, size and pore area in hundreds of images, where each individual image contains 15 to 60 stomata.

### Generalisation of SCAN application to other species

We collected images and conducted the manual-machine test on stomata density with six other species using the same devices and setup strategies as used for canola (Fig. 8). The results of manual-machine test on stomatal density indicates a higher correlation between SCAN and manual measurements, even for species outside our training domain (Fig. 8C). The manual-machine stomatal density tests were conducted on Arabidopsis, tobacco, rice, wheat, maize and proso millet. The corresponding correlation coefficients were 0.9582, 0.8919, 0.8123, 0.9834, 0.9106 and 0.8786, respectively. These high correlation coefficients indicates that SCAN can generalise well to other species, accurately predicting stomatal density even in species not included in the training set.

## Discussion

We developed a tool that provides rapid and reliable phenotyping measurement of canola leaf stomatal traits, including stomatal density, stomatal area, and pore area, by combining the use of high-resolution digital microscope and machine learning. We call this tool Stomatal Comprehensive Automated Neural Network or SCAN. Our model testing revealed that SCAN can produce consistent and reproducible predictions each time, eliminating variations in measurements introduced by human error during manual counting. In addition, we determined that SCAN could carry out the workflow of hundreds of images in minutes (Table 1), which is typically equivalent to more than one day’s workload for an individual researcher doing manual measurement. The use of high-resolution digital microscope in capturing leaf stomatal density and aperture status (i.e., open or closed) in SCAN also provided a non-destructive and automated analysis of stomatal features in real-time during phenotyping. The lightweight design of the digital microscope, straightforward pairing between the digital microscope and a laptop, and the offline accessibility of SCAN (Fig. 6) offer a portable and user-friendly tool for field application. The desktop application system of SCAN eliminates the technical burden from the users by packaging the three models in a user-friendly interface that enables processing of multiple images by just clicking the appropriate buttons in the display window (Fig. 6) – a major advantage of SCAN over other existing stomatal trait machine learning models. SCAN can also process the collected data into plots that can be exported along with the processed images and raw data file. SCAN can also potentially be correlated with other stomatal conductance tools (e.g., LI-600 Porometer /Fluorometer) to further study stomatal traits (Caine et al., 2023).

From SCAN predictions, we observed that the predictions hold at different magnifications (Fig. 8), even though only 700x magnification images of canola were used in our training set. We determined that SCAN is flexible enough in detecting stomata in different species of dicots and monocots (Figs. 7 and 8C).

However, the distinct guard cell shapes in dicots (kidney-shaped) and monocots (dumbbell-shaped) (Hetherington & Woodward, 2003) and different complex types of stomata (Lawson & Matthews, 2020) made it difficult for SCAN, which is specifically trained for canola, to accurately predict stomatal size and pore area in other species. In canola, the stomatal density predictions from SCAN are relatively more precise than that of stomatal size and pore area, based on the results from metrices evaluation (Fig. 4) and manual-machine tests (Fig. 7). From observations, the orientation of each stoma in canola is random, which can potentially decrease the accuracy of pixel-level segmentation. Moreover, human experts tend to measure the stomatal size or pore area as a maximum oval-shaped pattern based on the kidney-shaped guard cell morphology in dicot without zooming in, while SCAN calculates the area pixel by pixel. This difference in methodology causes the correlation to drop in stomatal area and pore area manual-machine tests (Fig. 7B-C). Difference in leaf structural features can also have influence on the quality of images obtained and hence model predictions. For instance, in Figure 8C, Arabidopsis showed lower accuracy in stomatal density predictions because abundant trichomes present on the Arabidopsis leaf surface create noise and blur in the background affecting the focus of the microscope images. Furthermore, the small size and inherent curvature in the leaf of the Arabidopsis plant made imaging fiddly. Prediction accuracy will improve for Arabidopsis with higher quality images or by retraining the model to tailor for Arabidopsis features, which can also be initiated using SCAN.

From the perspective of model capability, the potential of extending SCAN to other species offers two main benefits. First, SCAN’s weights can be used to train a new fine-tuned network for other species, which can reduce training time by transferring the pretrained network from SCAN. For instance, researchers can use SCAN as a starting point for measuring stomatal traits in other species based on the similar feature found in stomata. SCAN can then learn and conduct accurate prediction once new dataset of the species of interest is annotated and provided. This brings us to the second benefit, because SCAN can also serve as an annotation tool when preparing a new training dataset for other species, reducing the trivial and repetitive work in stomata and its corresponding pore labelling. In this case, researchers only need to annotate the remaining undetected instances in establishing the new dataset before retraining a new model for other species in measuring stomatal leaf traits.

## Conclusion

This study describes the development and application of the first automated leaf stomatal traits measurement tool for canola. Stomatal Comprehensive Automated Neural Network (SCAN) alleviates the statistical workload for biological researchers by providing accurate measurements of stomatal traits in high-throughput, portable, real-time and non-destructive manner. The user-friendly desktop application system allows researchers to focus on biological questions rather than the technical aspects of setup, operation, or data analysis issues in computer science. SCAN’s ability to produce consistent and reproducible results makes SCAN a valuable tool for phenotyping applications.

## Acknowledgements

We thank Ms. Amelia Desouza for her technical assistance and Dr. Abdeljalil El Habti for consultation and advice. This work was funded in part by The Australian Research Council Centre of Excellence for Translational Photosynthesis (CE140100015).

## Authors contribution

LY obtained the images, developed the machine learning models, and performed the analyses. SvC advised on the experimental design and data analysis. FRD conceived the project, designed and supervised the work, and analysed the data. All authors read and approved the final manuscript.

