## Supplemental information for "Automated and high throughput measurement of leaf stomatal traits in canola"

### Supplementary Tables

**Supplementary Table 1. Model templates and weights.**

This table details the model templates and transferred weights used in the different stages of SCAN's development. The stomata detection model is initiated using the small model template (yolov8s.yaml) from the YOLOv8 series and is trained from scratch. The stomata segmentation model uses the YOLOv8 segmentation template (yolov8s-seg.yaml) and inherits the transferred weights from detection model to continue training the stomata segmentation model. In addition, the pore segmentation model uses the YOLOv8 nano segmentation template (yolov8n-seg.yaml) and is trained on the Extracted Canola Stoma dataset with transferred weights from the stomata segmentation.

| Model | Model Template | Transferred Weights |
|---|---|---|
| Stomata Detection | yolov8s.yaml | N/A |
| Stomata Segmentation | yolov8s-seg.yaml | Stomata Detection Model |
| Pore Segmentation | yolov8n-seg.yaml | Stomata Segmentation Model |

**Supplementary Table 2. SCAN processing time in different processing unit.**

This table summarizes the SCAN processing time across three different processing units: a mid-range CPU (Intel(R) Core (TM) i7-6700 CPU @ 3.40GHz 3.41 GHz), a newer CPU (13th Gen Intel(R) Core (TM) i7-1365U 1.80 GHz) and a server GPU (NVIDIA A30 Tensor Core GPU 24G). Processing time was measured for the entire SCAN system workflow using two test datasets. The 132-image dataset is an unseen dataset containing canola microscopies of both abaxial and adaxial leaf surfaces, while the 427-image dataset was generated by selecting images from the training dataset. Each microscopy has resolution of 2560 x 1980. The table includes the total images processed and the processing time in seconds for each configuration, both with and without pore measurements.

| Process unit | Total images | Processing time in second with all measurements | Processing time in second without pore measurement |
|---|---|---|---|
| Intel(R) Core (TM) i7-6700 CPU @ 3.40GHz 3.41 GHz | 132 | 193.5 | 53.1 |
| | 427 | 548.9 | 122.7 |
| 13th Gen Intel(R) Core (TM) i7-1365U 1.80 GHz | 132 | 136.6 | 32.1 |
| | 427 | 253.1 | 75.9 |
| NVIDIA A30 Tensor Core GPU 24G | 132 | 50.9 | 10.5 |
| | 427 | 157.7 | 13.8 |

### Supplementary Figures

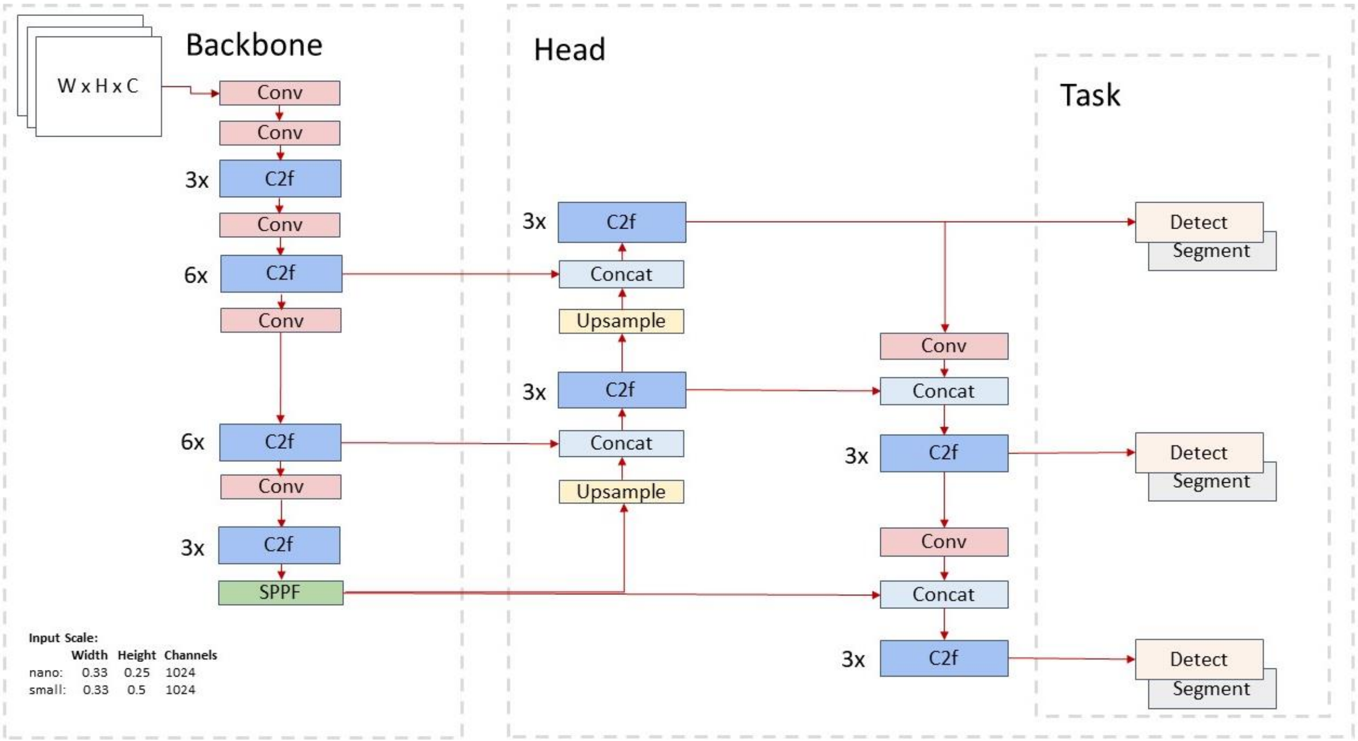


**Supplementary Figure 1. Architecture of YOLOv8 network used to develop SCAN.**

This figure summarizes the structure of YOLOv8 network structure.

**Input**: Microscope images with input scaling factor before connecting to backbone.

**Backbone:** CSPDarkNet53 (Wang et al., 2020) features extractor composed of convolution (conv), Cross Stage Partial Bottleneck with 2 convolutions-Fast(C2f), and Spatial Pyramid Pooling – Fast (SPPF) building blocks. C2f collects multi-level features before the next layer, and SPPF helps the model learn consistent features.

**Head:** Feature Pyramid Network (Lin et al., 2017) and Path Aggregation Network (Liu et al., 2018) (PAFPN) work together to deliver fast features fusion.

**Task:** The last layer is in head flexible to conduct bounding box localizing detection or pixel-level mask segmentation tasks.

**Algorithm 1:** Scale Bar Detection

**Input:** Image($image$), Scale bar Unit($unit$)
**Output:** Scale bar ratio from pixel to Unit($ratio$)
**Definitions:**

$roi$ - region of interests for scale bar location
$lines_p$ - filtered lines by $PolarCoordinate(\rho, \theta)$
$linesLengths_\alpha$ - lines lengths in $\alpha$-axis in $CartesianCoordinate(x, y)$
$ratio_{default}$ - default ratio value

**Try**
    $roi \leftarrow$ Bottom-Left($image$)
    $edges \leftarrow$ Canny Edge Detection($roi, \ldots$)
    $lines \leftarrow$ HoughLines Trasformation($edges, \ldots$)
    $lines_p \leftarrow lines$ where $\theta$ near 0 or 90 degrees
    $candidate_x \leftarrow \max(linesLengths_x)$
    $candidate_y \leftarrow \max(linesLengths_y)$
    $ratio \leftarrow unit/\max(candidate_x, candidate_y)$
    **return** $ratio$;
**Catch Exceptions**
    **return** $ratio_{default}$

**Supplementary Figure 2. Scale bar detection algorithm in SCAN.**

The algorithm aims to establish a bridge between pixel-level measurements and the scale bar unit for further analysis. It takes the original microscopy image and scale bar unit as inputs and return the corresponding ratio from pixel to the given unit. The algorithm effectively locates the scale bar in the bottom-left region of the image, applies edge detection and line transformation techniques, and calculates the ratio between the pixel length of the detected lines and the given unit. This ratio is used for converting pixel-level measurements to real-work units. The default ratio value will be safely returned if no line candidates present after or during the algorithm.
